# Transcriptomics Reveals Tumorigenesis and Pathogen Response in Scombrid Puffy Snout Syndrome

**DOI:** 10.1101/2023.06.26.546572

**Authors:** Savanah L. Leidholt, Kalia Bistolas, Manoj Pastey, Mark Dasenko, Emily Miller, Kyle S. Van Houtan, Andre Boustany, Tatiana Galivn, Rebecca Vega Thurber

**Author notes:** **Corresponding Author Information**: Mailing Address: 2820 SW Campus Way, Corvallis, OR 97331.

## Abstract

Scombrids represent some of the most economically important fisheries globally. However, increased interest in creating aquaculture systems for these fish has also increased the risk for disease emergence. One such disease, Puffy Snout Syndrome (PSS), causes collagenous tissue growths on the face in numerous scombrid taxa. PSS has mainly been documented in captive-held fish populations and can lead to high mortality rates. Despite this, little is known about the causative agent(s) of PSS and the immune response they elicit. Therefore, we leveraged transcriptomic data of PSS symptomatic-captive, asymptomatic-captive, and healthy-wild Pacific Mackerel (*Scomber japonicus*) to evaluate the physiological characteristics of PSS infections and identify a potential mechanism of disease. Captive symptomatic and asymptomatic mackerel showed distinct gene expression patterns from their wild counterparts. Genes involved in tumorigenesis, immune response, and tissue remodeling were overexpressed in captive-held fish. WNT9 was the most overexpressed gene in captive groups, and the WNT signaling pathway itself showed a ∼3 fold increase in KEGG pathway enrichment analysis in captive animals. When captive fish were compared, asymptomatic fish showed lower expression of inflammation genes, but high expression of tumor suppressor genes compared to symptomatic-captive fish. Together, these host pathophysiological data and our past visual identification of RNA virus-like particles in afflicted tissues suggest that viral-mediated oncogenesis may be driving PSS in captive mackerel.

## Introduction

The Scombridae fishes (tunas, bonitos, mackerel) comprise one of the largest global finfish economies, which generated $40.8 billion USD in 2018 alone [1]. In recent years, efforts to improve the efficiency of tuna ranching and farming have increased. This has involved keeping tunas captive for extended periods of time, potentially years, at high densities. Tunas and mackerels are typically hardy species in aquaculture with negligible economic loss from diseases [2–4]. However, high fish-stock densities, low-quality food sources, and increasing temperatures in mariculture operations have contributed to large-scale disease outbreaks [4,5]. Mass mortality outbreaks in tuna aquaculture are often associated with the parasitic blood fluke (*Cordicola* spp.). Mortalities have ranged from 50-80% of an enclosure [6]. Incidences of mass mortality events from bacterial and viral diseases are less common than parasitic pathogens, but significant die-offs occur in scombrid aquaculture, most often associated with juvenile fish. One prolonged disease outbreak in Spring 2004 in an Atlantic bluefin tuna, *Thunnus thynnus* operation resulted in 2500 fish deaths over a six-month period [3]. During the outbreak, 45 fish were sampled and 100% tested positive for a new isolate of *Photobacterium damselae* [3]. This outbreak was directly correlated with a sharp increase in water temperatures and compounded by the long-term feeding of low-quality bait feed [7]. There are also reports of mycobacterium infections in Atlantic bluefin tunas [8] and, more recently, in Atlantic mackerel, *Scomber scombrus* [9]. Other bacterial outbreaks reported in hatchery-reared Atlantic bluefin tuna larvae were caused by *Vibrio* and *Tenacibaculum* sp. [10]. Both nervous necrosis (NNV) and red seabream iridovirus (RSIV) have had high mortality rates in larval and juvenile Pacific Bluefin Tunas, but viral epizootics have yet to be described in adults [11–13]. However, one study described that juvenile Pacific Bluefin tuna were highly susceptible to RSIV when experimentally challenged via oral viral entry, with nearly 90% mortality in the infected group [14]. RSIV is also correlated with periods of elevated water temperature [11].

Puffy Snout Syndrome (PSS) is a common disease of captive scombrids characterized by an overgrowth of collagenous tissue on the fish face, which inhibits foraging. However, it remains unknown if this syndrome is caused by pathogen (bacteria, virus, parasite, etc), environmental challenge, or a genetic predisposition. This syndrome has been reported in several species of scombrid fish including: yellowfin tuna (*T. albacares*), Atlantic and Pacific bluefin tuna (*T. thynnus)*, blackfin tuna (*T. atlanticus)*, skipjack tuna (*Katsuwonus pelamis)*, kawakaw (*Euthynnus affinis*), Pacific bonito (*Sarda chiliensis*), Pacific mackerel (*S. japonicas)*, and Atlantic mackerel (*S. scombrus)* [15–21]. Outside of the scombrid fish family, the disease has been seen in northern anchovies (*Engraulis mordax*), Pacific sardines (*Sardinops sagax)*, rainbow trout (*Onchornycus mykiss)*, California yellowtail (*Seriola dorsalis*), and barracuda (*Sphyraena sp.*) [21,22].

In tunas and mackerel, the first signs of PSS are a darkening and wrinkling of the skin, though this phenotype is more pronounced in tunas. As the disease progresses, a gelatinous substance overgrows the dorsal area of the head, often occluding the eyes. Work using Masson trichrome staining found that normal facial tissue was composed primarily of striated muscle fiber; however, in PSS-afflicted fish, this tissue began to break down and become replaced by collagen [21]. Further, mucus-producing goblet cells, important for creating a healthy slime coat on the fish, were noticeably decreased or malformed in PSS-affected tunas [21]. Severe degradation at the cellular level renders mitochondria as one of the only recognizable structures in severe PSS tissues [22]. A seminal survey of 11 facilities that reared or held tunas, mackerels, or both, found that of the total 22 rearing units, PSS developed in 68% of these units [21]. Typically, fish with severe PSS are unable to feed and, as a result, are either euthanized or starve to death. Further, PSS has only been observed in captive fish, though this may be due to survivorship bias. None of the facilities surveyed reported seeing PSS in wild-caught fish, with PSS eventually developing in the first six months of captivity [21].

Despite its unique disease characteristics, density-dependent prevalence, and wide host range, PSS remains understudied but highly important, as severe PSS has mortality rates ranging from 20-100% of an enclosure [15–17,21]. Further, PSS appears to be species-specific as reports describe other fish taxa, such as Mahi Mahi (*Coryphaena hippurus*), in the same enclosures with PSS-afflicted scombrids, but without PSS signs [21]. Only 9 published reports exist on the topic, and only two are peer-reviewed and attempt to identify etiological agents [21,22]. To understand the etiological agents potentially responsible for PSS, our recent work used electron microscopy techniques and identified viral-like particles (VLPs) within PSS-afflicted tissues of the Pacific mackerel (*S. japonicus*) [22]. These VLPs shared morphology with many small RNA viruses being amorphous in shape and ∼100 nm in diameter. Along with VLPs, a high abundance of deformed mitochondria were identified in these same PSS-afflicted tissues. The VLPs and disease prevalence in increased fish stock densities [21] strongly suggest that PSS onset or severity is associated with a viral infectious agent. Interestingly, the number of malformed mitochondria within both asymptomatic-captive animals (without visible signs of disease) and symptomatic-captive animals (with visible signs of disease) was the same suggesting that some tissue level signs of PSS may begin to occur even in visually healthy fish.

As a next step to identify the underlying mechanisms of PSS, we leveraged high throughput sequencing methods to further characterize this disease at the physiological and genomic levels. Our aims were to 1) analyze comparative host gene expression dynamics in PSS of apparently healthy mackerel, asymptomatic-captive, and symptomatic-captive mackerel; 2) identify expression pathways indicative of specific diseases (i.e. viral, bacterial, cancerous). Here we present the results of differential gene expression analysis for *S. japonicus* afflicted with PSS and, for the first time, evidence of tumorigenesis in this species.

## Methods and Materials

### Sample Collection

Sample collection and euthanasia of captive-held and wild-caught mackerel were performed as described previously [22]. Briefly, captive-held Pacific mackerel were obtained from the Monterey Bay Aquarium (MBA) during a routine deaccession and euthanized with an overdose of MS-222 (i.e. tricaine mesylate). Wild-caught fish were captured and euthanized on site from the coast of Monterey Bay, California, USA. All fish were placed in separate bags and placed on ice or in -20_ until facial necropsy. Epithelial and muscle tissue were taken from behind the nares and anterior to the eyes. A total of 3 symptomatic-captive, 3 asymptomatic-captive, and 3 wild-caught control (i.e., wild control mackerel) were used for this analysis. These samples were a subset of fish used in Miller et. al. 2021.

### RNA Extraction

Frozen mackerel samples were stored in a -80□ freezer and held in liquid nitrogen before RNA extraction to ensure RNA stability within tissues. To reduce the possibility of cross-contamination, all surfaces were sterilized with 95% ethanol, followed by bleach. Gloves were exchanged between each sample to avoid cross-contamination. Following aseptic protocols, approximately 0.5 mg of tissue was removed with a sterile scalpel and forceps, homogenized in liquid nitrogen using an autoclaved mortar and pestle, and extracted using the E.Z.N.A DNA/RNA Isolation kit from Omega Bio-tek per manufacturer instructions [23]. RNA and DNA were eluted in 50 µl of nuclease-free water and stored at - 80□ until library preparation.

### Meta-transcriptome Library Preparation

RNAseq library preparation and sequencing were performed at the Center for Quantitative Life Sciences at Oregon State University. Briefly, ribosomal RNA was removed from the total RNA extractions using an Illumina mouse/rat/human Ribo-Zero kit for host rRNA removal and then a second Ribo-Zero kit for bacterial rRNA removal. After rRNA removal, a stranded RNA-seq library was prepared using the Takara PrepX DNA library prep. Then samples were sequenced on one lane of a 2x150bp PE Illumina HiSeq 3000 run.

### Sequence Alignment and Expression Analysis

Illumina HiSeq libraries ranged from 28,745,252 to 49,799,840 paired-end reads. The sequence data was first quality-filtered by removing reads with a Phred score lower than 30 to demonstrate 99.99% of sequencing base call accuracy. Adapters and poly(A) tails were removed from the reads and normalized to 30x coverage using the BBtools suite [24]. Library read counts after each quality control step can be found in Supplementary Table 1. The resulting 593,829,134 reads were then merged using BBmerge [25], and the resulting files were used to construct a *de novo* transcriptome of *Scomber japonicus.* Trinity was used to build the transcriptome using default settings (v3.9.1; [26]). GC content of the *de novo* transcriptome was 41%. TransDecoder (v5.5;[27]) was used to identify candidate coding regions within the *de novo* transcriptome. These coding regions were annotated using a BLASTx database search against the KEGG protein database (v108.0;[28]). Quality-controlled transcripts of individual libraries (i.e. samples) were mapped to the *de novo* assembled transcriptome using the Salmon Estimation Method packaged with Trinity (v.1.13.0;[29]). Compared to two additional reference-guided assemblies (*S. colias* genome and *S. scombrus* transcriptome), the *de novo* transcriptome recruited the most reads among libraries and was therefore used in this study. All three assemblies’ metrics and recruitment statistics can be found in Supplemental Material 1. Individual read libraries are available at NCBI under BioProject PRJNA967399.

After removing outliers, DESeq2 (v.3.16;[30]) was employed with R statistical programming software (v.4.1.2;[31]) to test for differential expression between pairs of treatment groups to determine whether each model coefficient differed significantly from zero. Sequences with adjusted p-values less than 0.05 and an absolute value of log2 fold-change greater than 2 were defined as differentially expressed genes (DEGs). A permanova was then used through the VEGAN (v2.7; [32]) package in R statistical programming software to detect differences amongst the three groups over 999 iterations. The KEGG mapper reconstruct tool (v5.0; [33]) was used to group genes into global overview pathways. ShinyGo (v0.77; [34]) was used to perform KEGG pathway enrichment analysis using the closest related species *Danio reiro* for the reference as has been done previously with other non-model teleost species [35–37]. ShinyGo v0.77 was set to remove duplicate IDs with all other parameters as default.

## Results

In this study, we used comparative transcriptomics to evaluate physiological variation in Pacific mackerel (S. japonicus), including wild fish and captive fish with visible signs (symptomatic) and without visible signs (asymptomatic) of Puffy Snout Syndrome (PSS, Fig 1A). Wild-caught mackerel were assumed to be healthy in all cases as there were no visible signs of disease externally or internally. DEG analysis, plotted in principal component space and tested using a Permanova with 999 iterations (p-value =0.008), revealed that captive-status accounted for the most variability within gene expression amongst our groups, explaining 25% of the variance (Fig 1B).

**Fig 1.**
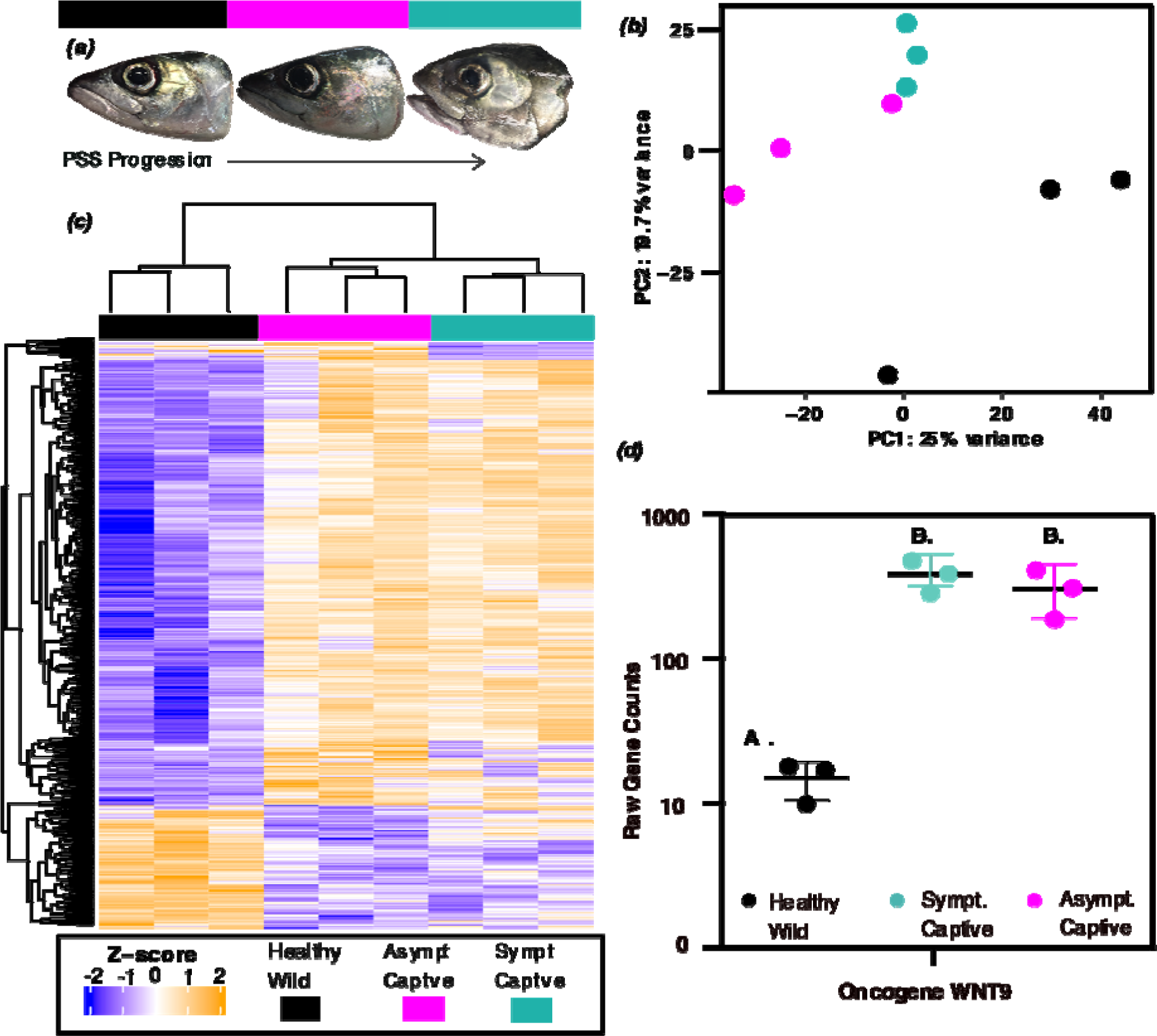
Transcriptomic data shows distinct differences between captive and wild mackerel. A) Adapted from Miller et al. 2021. From left to right images of healthy wild fish, asymptomatic captive fish, and PSS symptomatic captive fish. B) Principal component analysis (PCA) plot of all transcripts showing the top 2 axes which explain the most variance. A permanova was performed on the PCA and found that variances among groups were significantly different (p-value = 0.008). C) Differentially expressed genes in a heatmap showed clear differentiation by fish captive status as seen in the hierarchical clustering. D) Raw counts of the top differentially expressed gene, *Wnt-9B* in each group. Black boxe and symbols represent healthy wild mackerel, green represents symptomatic captive mackerel, and pink represents asymptomatic captive mackerel.

### Gene Expression and KEGG Enrichment Analysis Identified Three Overarching Gene Groups

A total of 1,115 DEGs were identified between all pairwise comparisons as containing large coding sequences and characterized in the KEGG database (Fig 1C). Initially, DeSeq2 identified a total of 2,155 differentially expressed genes (DEGs) between asymptomatic-captive and wild-control groups. However, of these, 806 were classified as coding sequences by TransDecoder with open reading frames (ORFs) larger than 100 amino acids, but only 567 of these genes were characterized by the KEGG database. Between symptomatic-captive and wild-control groups, 4,183 DEGs were found of which 793 contained ORFs larger than 100 amino acids and 526 were characterized within the KEGG database. Lastly, Deseq2 identified 85 DEGs between asymptomatic-captive and symptomatic-captive mackerel with 35 of these containing long ORFs and 22 of these characterized within KEGG (Supplemental table 2). Though a large portion of sequences were removed with this approach, this method allowed for conservative downstream analysis and annotation of DEGs. To determine which genes might be diagnostic of PSS, we also identified DEGs that were highly expressed in the symptomatic fish cohort. Wingless-related integration site 9 (*wnt9*) was identified as the most highly differentially expressed gene within symptomatic-captive-held *S. japonicus* (p-value = 1.31^-35^) (Fig 1D). Surprisingly, this gene was also the 2^nd^ most expressed gene in asymptomatic-captive tissue compared to wild healthy mackerel (p-value = 1.38^-30^).

Once annotated, genes for each pairwise comparisons were used for KEGG enrichment analysis among the fish cohorts using ShinyGO to identify differences and commonalities between wild fish and each of the captive fish health states as well as between the asymptomatic and symptomatic fish. Between asymptomatic-captive and wild-control groups, 79% of genes matched to reference pathways. Similarly, 79% of genes matched to reference pathways between symptomatic-captive and wild-control, and 68% of genes matched to reference pathways between asymptomatic and symptomatic captive (Fig 2). All genes mapping to pathways can be found in supplementary table 3. Significantly enriched pathways and DEGs were broadly categorized into three groups including: 1) tumor-related, 2) immune-related, and 3) pathophysiological response to tissue degradation genes and pathways. However, some of the enriched pathways and individual differentially expressed genes can be broadly characterized into more than one of these defined groups.

**Fig 2.**
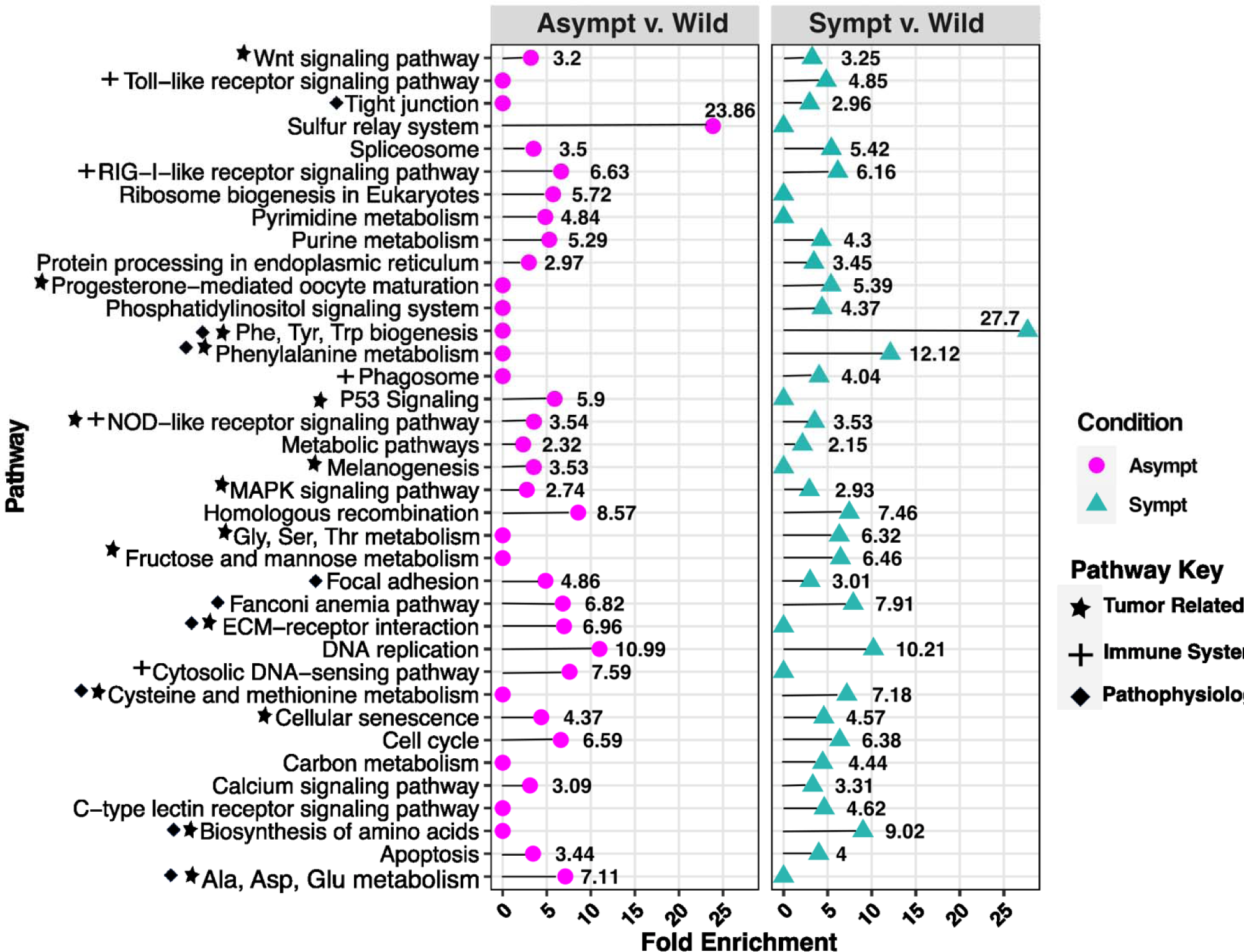
KEGG pathway enrichment analysis (FDR p. adjust < 0.05). Enriched pathways between asymptomatic-captive (pink) and healthy-wild groups shown in the left panel. Enriched pathways between symptomatic-captive (green) and healthy-wild groups based on DEGs. Stars represent pathways that are related to tumorigenesis or suppression, crosses represent pathways of the immune system, and diamonds show pathways relating to pathophysiology. Some pathways fit into multiple of these categories.

### Captive fish share more similar transcriptomic profiles compared to wild-control fish

Tumorigenesis genes and pathways were similarly enriched in captive fish compared to wild fish regardless of disease status. Pathways enriched between the asymptomatic-captive and the wild-control group that promote tumorigenesis were the mitogen-activated protein kinase pathway (*MAPK*) with 2.73 fold enrichment, wingless integration site pathway (*WNT*) with 3.19 fold enrichment, calcium signaling pathway with 3.09 fold enrichment, and melanogenesis pathway with 3.53 fold enrichment (Fig 2). Tumor protein 53 (*TP53*), a tumor suppressor pathway, was also enriched between these groups by 5.89 fold (Fig 2). Tumorigenesis pathways enriched between symptomatic-captive and wild-control groups were *MAPK* (2.93 fold enriched), *WNT* (3.24 fold enriched), and calcium signaling pathway (3.31 fold enriched) (Fig 2). No tumor suppressor pathways were enriched between symptomatic-captive and wild-control groups.

Other pathways indirectly related to tumorigenesis and enriched in both captive groups were amino acid biogenesis and metabolism, nucleic acid metabolism, and sugar metabolism. Genes identified as prognostic markers to human cancers and overexpressed in captive fish tissue include: *WNT9*, *WNT7a*, *WNT3*, cyclin E (*CCNE2*), low-density lipoprotein (*LDL*), latent transforming growth factor beta binding protein 1 (*LTBP1*), retinoblastoma protein 1 (*RB1*), and baculoviral IAP repeat-containing protein 5 (*BIRC5*) (Fig 3A).

**Fig 3.**
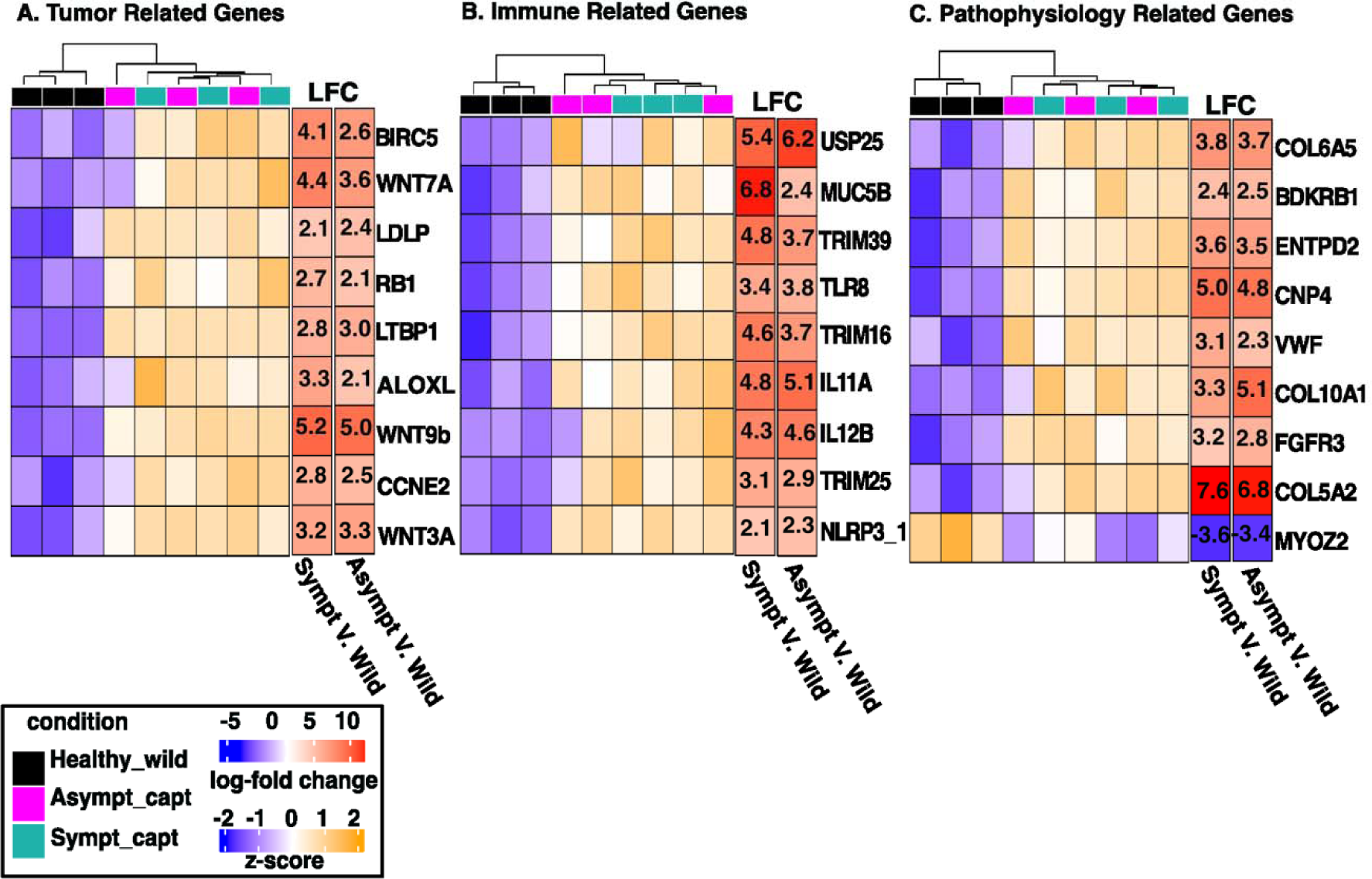
Heatmaps of three groups of differentially regulated genes. Selected A) tumor-related; B) immune response; C) and pathophysiological marker genes. Black boxes and symbols represent healthy wild mackerel, green represents symptomatic captive mackerel, and pink represents asymptomatic captive mackerel. Smaller heatmaps to the right of each main graph are the log2 fold-change values for the symptomatic and asymptomatic groups, respectively.

The Innate immune system pathways also showed high enrichment in captive fish compared to wild-control fish (Fig 2) and were linked with pathways such as: inflammation, viral immune response, and cancer formation. Immune pathways enriched in captive fish were interestingly affiliated with pathogen infection such as elevation of pattern recognition receptors (PRRs). PRR pathways enriched between asymptomatic-captive fish and wild-control fish include nucleotide-binding oligomerization domain-like (NOD-like) receptor pathway (3.54 fold), retinoic acid-inducible gene (RIG-1) receptor pathway (6.63 fold), and the cytosolic DNA sensing pathway (7.6 fold) (Fig 2). Similarly, PRR pathways were enriched between symptomatic-captive fish and wild-control fish include the NOD-like receptor pathway (3.54 fold), RIG-1 receptor pathway (6.16 fold), and the toll-like receptor (TLR) signaling pathway (4.85 fold) (Fig 2).

Innate immune system genes showed high overexpression in captive fish compared to wild-control fish (Fig 2). The PRR NLR family pyrin domain containing 3 transcript version 1 (*NLRP3_1*) and toll-like receptor 8 (*TLR8)* show ∼2 and 3 fold increase in captive fish respectively. Tripartite motif-containing (*TRIM*) proteins, *TRIM16*, *TRIM25*, and *TRIM39* were overexpressed in captive fish compared to wild by ∼2, 4, and 4 fold respectively. General inflammation response genes overexpressed in captive fish included: interleukin 11 (*il-11a*) with ∼5 fold increase, interleukin 12b (*il-12b*) with ∼4.5 fold increase, ubiquitin-specific protease 25 (USP25) with ∼6 fold increase, and mucin 5 (*Muc5a*) with ∼2-6 fold increase (Fig 3B).

Enriched pathways and genes associated with tissue and cellular repair in response to degradation were prominent in captive groups. The extracellular matrix receptor (ECM) pathway, focal adhesion pathway, and fanconi anemia pathway were enriched by 6.96, 4.86, and 6.82 fold in asymptomatic-captive fish compared to wild-control fish (Fig 2). Symptomatic-captive fish were enriched in the latter two pathways by 3 and 7.91 fold compared to wild-control fish (Fig 2).

Genes related to tissue and cellular regeneration were overexpressed between 1.5-7 fold increase within captive-held mackerel (Fig 3C). The most abundant of these genes induces collagen production (*Col5A2, Col10a1, Col6a*). Genes associated with bone development and maintenance (fibroblast growth factor receptor 3; *FGFR3*), fibrosis (Ectonucleoside Triphosphate Diphosphohydrolase 2; *ENTPD2*), and vascular repair (Bradykinin Receptor 2; *BDKRB2*) were also overexpressed in captive-held mackerel. Myozenin 2 (*MYOZ2),* a gene responsible for skeletal muscle formation and growth is significantly underexpressed in both of our captive groups compared to the wild by ∼ 3.5 fold. Tissue overgrowth can also be associated with tumorigenesis (collagen genes and FGF3) or a general response to inflammation Von Willebrand Factor (*VWF*) (Fig 3C).

### Tumor suppressor and inflammation genes were differentially expressed between asymptomatic and symptomatic captive fish groups

Since the initial aim of the project was to explore mechanisms of disease in PSS fish, and because we found that captive fish had more differences to wild than expected, we then compared DEGs only between the two captive fish cohorts, symptomatic and asymptomatic. Extracellular matrix (ECM) signaling was the only pathway significantly enriched between captive groups with a 78 fold enrichment in asymptomatic fish (Fig 4A). An investigation into the DEGs between captive groups showed high expression of inflammation related genes in symptomatic fish: platelet and endothelial cell adhesion molecule 1 (*PECAM1*) with 3 fold overexpression, *NLRP3_2* with 5-fold overexpression, and arginine vasopressin receptor 1 (*AVPR1B*) with 2.5 fold overexpression (Fig 4B). Genes responsible for tumor suppression were significantly underexpressed in symptomatic-captive fish compared to asymptomatic-captive fish. Tumor suppressor genes of note are Deleted in malignant brain tumor 1 (*DMBT1*) and sterile α motif and HD domain-containing protein 1 (*SAMHD1*). These genes were under-expressed in symptomatic fish by 7.74 and 7.72 fold, respectively (Fig 4B).

**Fig 4.**
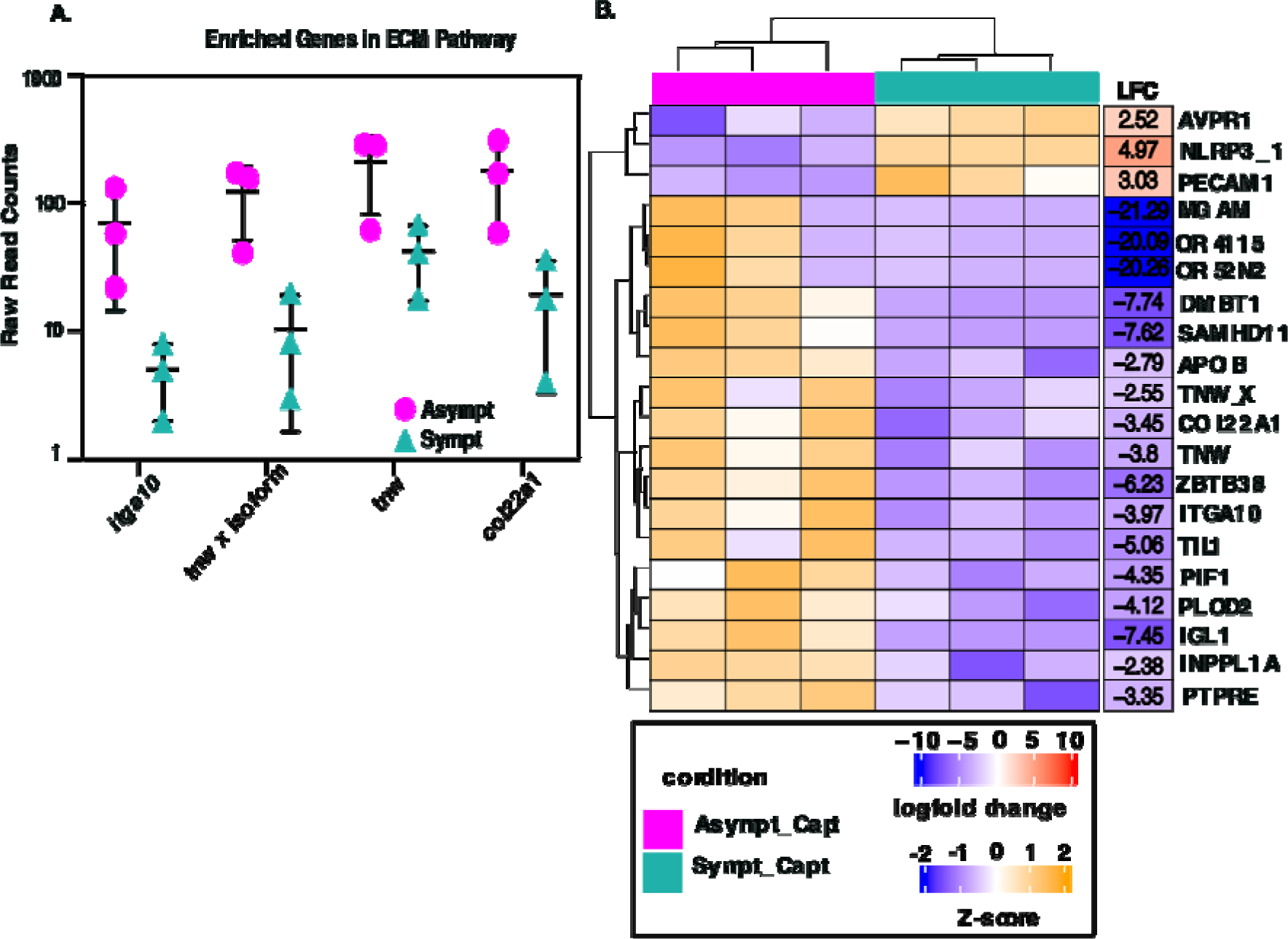
Transcriptional differences between captive symptomatic and asymptomatic fish. A) KEGG enrichment analysis of DEGs between captive groups (FDR p. adjust < 0.05) B) Heatmap showing DEGs between captive groups. Smaller heatmap to the right represents the log-fold change (LFC) of each gene between asymptomatic and symptomatic captive mackerel. Pink boxes represent asymptomatic captive mackerel, and green boxes represent symptomatic captive mackerel.

## Discussion

This work characterized Puffy Snout Syndrome (PSS) at the transcriptomic level, showing evidence of facial tumorigenesis through the overexpression of pan-cancer markers within the damaged tissues and potential avenues for infection. However, PSS remains an enigmatic syndrome with multiple focal areas for future study. Fish have both innate and adaptive immune responses to pathogens, yet tend to rely more heavily on the innate system, with key innate immune genes activated when presented with stress or microbial challenge. In this study, transcriptomics offered insight into variations in PSS physiology by identifying quantitative differences in gene expression amongst healthy, asymptomatic, and symptomatic fish. Here, we found that expression of tumorigenesis, immune response, and pathophysiology genes were dramatically divergent in captive fish compared to their wild counterparts. Transcriptional differences between the two captive groups showed DEGs related to poor survival prognosis in cancers and increased inflammation; these shifts likely play intricate roles in PSS related tumorigenesis, tumor suppression and gross pathology.

### Captive fish tissues are highly enriched in genes involved in tumorigenesis

In this study our transcriptomic data suggested that the primary presentation of PSS is the result of cell proliferation dysregulation and differentiation, likely leading to the observed collagenous growths and progression through hyperplasia and dysplasia in facial tissues. This process is often characteristic of cancers. The association of PSS with cancer formation is reflected by the numerous pathways found to be enriched in tumor related processes, such as *MAPK* and *WNT*. The mitogen-activated protein kinase (*MAPK*) pathway is pivotal to the regulation for gene expression, cellular growth and cell survival with malfunctions in this pathway resulting in uncontrolled cellular growth and division, often leading to melanomas [38]). MAPK can be stimulated by the gene WNT3a which is also overexpressed in captive fish. Similarly, the wingless integration site pathway (*WNT*) is involved in early cell development, cell differentiation/migration, and tissue homeostasis in adult vertebrates [39]. As mentioned previously, the gene *Wnt9b* was one of the top differentially expressed genes in captive animals representing the largest shift in gene expression. *Wnt9b* is the starting component of the canonical *WNT* pathway whose genes are often diagnostic for many human and non-human animal cancers [40–42]. In *Wnt9b* gene KO mice, facial defects have been observed [43,44]. It has been shown that WNT3, WNT9b and R-spondin (RSPO) integrate with other WNT signaling regulators to generate fine-tuned WNT/β-catenin signaling during facial morphogenesis [45]. This cooperation likely regulates cell proliferation through Fibroblast growth factor (FGF) signaling in the facial processes [45]. This suggests that overgrowth of collagenous tissue on the fish face in PSS may be because of overexpression of MAPK and WNT pathways.The overproduction of cell growth, survival, and proliferation led by *MAPK* and the *WNT* pathways in the tumor microenvironment is a metabolically costly endeavor. Tumors often show enrichment in biosynthetic activities such as amino acid and sugar metabolism to support the cell ’s overproduction of DNA replication, cell division, and tissue growth [46]. Our results are consistent with this observation, including pathways enriched in amino acid biosynthesis, nucleotide biosynthesis, and sugar metabolism in captive fish.

In aquaria held fish, tumor suppressor pathways were under-expressed in symptomatic fish compared to asymptomatic, suggesting that it is a hallmark of the PSS phenotype.. The tumor protein p53 pathway (P53) regulates the genes involved in DNA repair, apoptosis, cell cycle, and differentiation and has long been known as a tumor suppressor pathway [47,48]. Dysregulation in the P53 pathway can elicit cancer formation through pathway inhibition from oncoproteins or by gene mutations. These P53 gene pathways were underexpressed in captive symptomatic fish [49]. Similarly, retinoblastoma protein (*Rb1)* was significantly overexpressed in both captive fish groups. *Rb1* is a tumor suppressor gene that regulates mitosis and gene expression by binding chromatin and limiting gene expression of proteins involved in moving cells from the G1 to the S stage [50]. However, phosphorylation of *Rb1* by the cyclin-mediated pathway alters its ability to bind these genes [51]. CCNA2 encodes cyclin-A2, which regulates this transition, and is also highly expressed ∼1.5 fold (S1 table) suggesting that tumor suppression activity of Rb1 may be inhibited upstream by the overexpression of cyclins.

### Pathophysiology and cellular repair in response to degradation or to tumor formation in PSS

Captive fish exhibited enriched pathways and genes consistent with tissue repair and with tumor formation. The extracellular matrix receptor (ECM) pathway’s functional role is in tissue and organ structure maintenance and morphogenesis and was enriched in captive animals (Fig 2). The ECM is composed of a complex network of macromolecules that surround cells and tissues to provide structural support and aids in intracellular signaling [52,53]. The ECM pathway can be transcriptionally enriched in human cancers and is hypothesized to cloak cancerous cells from the immune system [54]. For example, the focal adhesion pathway (within the ECM pathway) is responsible for linking the actin skeleton to the ECM and serves a role in cell proliferation, growth, and migration. Voorhees, 2015 reported an initial fibrogenesis network formation in early onset PSS which was confirmed using Masson trichrome staining where PSS afflicted individuals showed a decrease in muscle content and increase in collagen within PSS tissues. This is consistent with findings in human melanomas where tenascin and fibronectin genes alter the collagen formation within the ECM, resulting in more rigid tumor formation [55]. A total of four different collagen types were found to be overexpressed in our PSS captive animals. Collagens have been described as a double edge sword of the tumor microenvironment where they not only drive immune invasion but also promote oncogenesis by releasing biochemical signals among cancer cells [56].

We also identified substantial evidence of tissue remodeling and wound healing in PSS animals based on the elevation of genes involved in vasodilation (*BDKRB2*), platelet binding (*VWF*), and bone regeneration (*FGFR3*). The observed altered gene expression profiles may explain some of the observed gross and cellular morphology of PSS, such as presence of increased tissue growth on the head and general increase in tissue inflammation. Further this alteration of tissue remodeling and wound healing is consistent with the Miller et al., 2021 findings where visually asymptomatic fish showed internal signs of extensive tissue degradation and higher percentages of malformed mitochondria along with their symptomatic counterparts in comparison to wild fish.

### Infection and stress may explain variation in the expression of immunity genes in captive fish

We hypothesize that altered immune-specific transcriptomic profiles in captive fish may result from prolonged physiological stress in captivity or an underlying pathogenic infection such as a virus or bacteria that captivity may exacerbate. Shifts in essential immune pathway gene expression are often indicative of animals responding to pathogens. In this study, we also found extensive changes in immune gene expression profiles. Three types of pattern recognition receptor (PRR) pathways such as Toll-like receptors (TLRs), nucleotide oligomerization domain (NOD)-like receptors (NLRs), retinoic acid-inducible gene-I (RIG-I)-like receptors (RLRs), were enriched in captive fish compared to wild (Fig 2). Some TLRs (TLR1, 2, 4, 5, 6, 10) are expressed on the surface of immune cells, mainly recognizing the membrane components of pathogenic microorganisms, such as lipids, lipoproteins, and proteins [57,58]; others (TLR3, 7, 8, 9) are expressed in the cytosol, which mainly recognize the nucleic acids of microorganisms [58]. NLRs are intracellular PRRs, which play a key role in sensing molecules associated with intracellular infection and stress. RLRs are a type of intracellular PRRs which can sense viral RNA molecules and play essential roles in innate antiviral immunity [59]. The DNA sensing pathways target DNA viruses or bacteria specifically in the cell cytosol.

Several DEGs we identified to be altered in captive fish are known to be responsible for host defense against viruses in teleosts. For example, tripartite motif 16 (*TRIM16*) is part of the fish-specific TRIM proteins and has been associated with viral infections in fish through negatively regulating interferon production [60–63]. Higher expression of TRIM16 is associated with higher viral loads and poor survival outcomes of fish [61]. Critically, TRIM25 has been one of the most commonly detected proteins in transcriptomic studies of virus infected fish or fish tissues in numerous species [64–67]. TRIM25 activates retinoic acid inducible (RIG-I) gene signaling by ligating polyubiquitin chains on the N terminal CARD [68]. This in turn induces downstream signaling by the RIG-I pathways. TRIM39 also referred to as bloodthirsty related trim 39 (*btr39*) is found to regulate the cell cycle during iridovirus and nodavirus infection in grouper (Epinephelus spp.) [69]. However, it exhibits less specificity and is induced upon both virus and bacterial challenge [69]. Further, toll-like receptor 8 (*TLR8*) in fish primarily functions to identify single-stranded RNA molecules and is presumed as a sensor for single-stranded RNA viral infection [70].

The inflammasome is a complex formed by the innate immune system in response to pathogen challenge, traumas, or chemical damage and have emerged as important regulators of cancer development and control [71]. The inflammasome induces host inflammation through activation of numerous cytokines. In teleosts, only NACHT-leucine-rich-repeat protein-1 (NALP1) and NLR family pyrin domain containing 3 (NLRP3) are capable of forming a complete inflammasome complex [72,73]. NLRP3 formation has been triggered in fish by bacterial LPS [72,74], cellular oxidative stress [75], and exposure to toxic metals [76]. Mitochondrial damage and dysfunction also have been observed in direct relation to the signaling pathways for NLRP3 induction [77,78]. We previously identified mitochondrial malformation in captive held animals. This alteration to NLRP3 may be directly or indirectly responsible for the mitochondrial damage that is consistent with our electron microscopy imaging results. In Miller et al. 2021 we showed PSS symptomatic fish had severe degradation and malformation of mitochondria in PSS afflicted tissues. Unlike the TRIM family proteins, however, it remains unclear whether the inflammasome complex is responding to pathogen challenge or environmental conditions in PSS affected mackerel. Further, inflammation is the hallmark of cancer development in animals [79], with the inflammasome playing a major role in the progression of the tumor environment. The role of the NLRP3 inflammasome in cancer promotion is often mediated by IL-1β, which is produced by various cells in the tumor, including tumor-associated macrophages (TAMs) and cancer-associated fibroblasts (CAFs) [80]. The products of the inflammasome NLRP3 -the pro-inflammatory cytokines-have been found to have opposing roles in different cancer types. Primarily in colorectal associated cancers NLRP3 has been identified as a tumor suppressor [81], however, the cytokines produced by NLRP3 are pro-tumorigenic in inflammatory induced gastric and skin cancers [82,83].

### Asymptomatic-captive fish likely represent an intermediate stage of the PSS disease

Comparison of the two captive groups revealed that symptomatic-captive fish showed higher expression of inflammation related genes and exhibit lower expression of tumor suppressor genes relative to asymptomatic fish (Fig 4b). These tumor suppressors are critical for regulating cell growth and proliferation and thus are often repressed at the transcriptional level in malignant tumors. This observation suggests that the asymptomatic captive fish with tumor suppressor gene expression may not exhibit initial signs of PSS because they can still transcriptionally maintain cell homeostasis.

For example, sterile α motif and HD domain-containing protein 1 (*SAMHD1*) is a tumor suppressor which was underexpressed in symptomatic fish compared to asymptomatic (Fig 4B). *SAMHD1* functions as a tumor suppressor by negatively regulating the innate immune response/inflammation, repairing DNA damage, and aiding in completing DNA replication by resolving stalled replication forks [84–86]. *SAMHD1* is underexpressed in several cancer types in humans and mice [59,84,87] though the mechanisms for each cancer are not yet fully understood.

Similarly, Deleted in malignant brain tumors 1 (*DMBT1*), is considered a tumor suppressor gene because it is a component of the mucosal immune system responsible for binding pathogens [88–90]. *DMBT1* represses cell proliferation, cell migration, and the invasion of cancer cells. *DMBT1* exhibited the lowest expression in wild-caught mackerel and symptomatic fish (S2 Table). In humans, this gene was underexpressed in skin and among other cancers [91–96], but overexpressed in other tumors [97–99]. Our results for *DMBT1* gene underexpression are consistent with other skin cancer transcriptomics and further support the hypothesis that asymptomatic-captive fish are at an intermediate disease state.

## Conclusion

We used RNA sequencing on three groups of pacific mackerel (wild, asymptomatic, and symptomatic) to pair with our previously published work on electron microscopy (EM) imaging of Puffy Snout Syndrome (PSS) [22]. Previous microscopy showed no evidence of bacterial infection, but instead identified small viral particles and large numbers of deformed mitochondria in afflicted fish with PSS. Aquaria fish with collagenous growth on the head were considered diseased (symptomatic with PSS). Conversely, wild fish were assumed to be healthy in all cases and those in aquaria presenting altered transcriptional profiles but without signs of disease were considered “asymptomatic”. The most significant observed changes in gene expression were detected between wild-caught fish and captive fish with genes involved in tumorigenesis pathways being the most elevated gene category in captive fish. Based on this combined transcriptomic evidence and visual disease phenotype, we propose that PSS is a condition which causes the formation of facial tumors in captive fish via alteration of the cell cycle and cell differentiation. Why these changes occur in captivity is currently unknown.

However, we also found significant shifts in tumor suppressor and inflammation/immune response genes between the two captive groups. We predict that PSS may be induced by an infectious oncogenic virus that induces tumorigenesis, a non-infectious physiological cancer, or by infectious cancer cells (transmissible cancer) not previously described in wild fish. Nonpathogenic transmissible cancers are extremely rare in wildlife, with few exceptions such as in marine molluscs, terrestrial canines, and rodents, and we can likely exclude this option. Given the visual observations from electron microscopy and increased expression of tumorigenesis genes and those involved in the innate immune response, a viral infection-inducing tumorigenesis may be the most probable candidate. However, although we cannot rule out the other hypotheses of physiological or transmissible cancers, we aim to further characterize the presence and abundance of viral-specific genes and genomes to pursue the oncogenic virus hypothesis in the future.

## Acknowledgements

This study was made possible by the generous contributions of the Monterrey Bay Aquarium, who provided us with the samples for this study. Specifically, we would like to extend gratitude to the husbandry operations team and animal care staff, who cared for and maintained all the animals used in this study.

S2_Table1: Read totals after each cleaning and normalization step. R1 represents the forward read pairs and R2 the reverse read pairs.

S3_Table2: Differentially expressed genes confirmed as coding sequences.

The first column describes the pairwise comparison. Only sequences that were found to be differentially expressed between a comparison, containing an open reading frame longer than 100 bp, and identified within the KEGG protein database were kept for downstream analysis.

S4_Table3: Total gene descriptions of annotated DEGs between all pairwise comparisons of fish groups

S5_Table4: KEGG Enrichment gene mapping ids between asymptomatic and wild DGEs

S6_Table5: Assembly information of the *De Novo* Pacific Mackerel transcriptome. General assembly statistics are in the right-most column.

---

**S1_Figure1:**
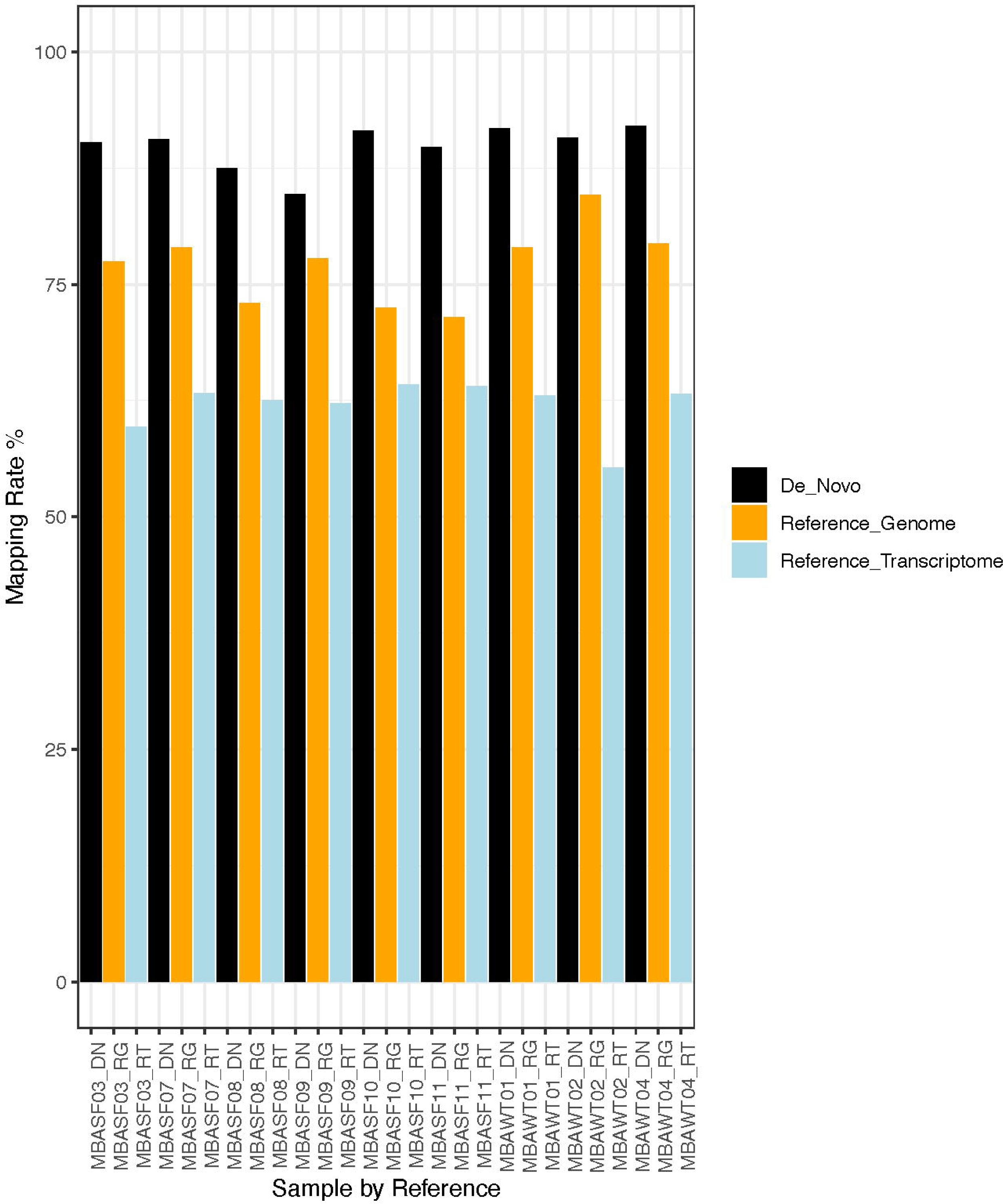
Mapping rate of individual reads to the *S. colias* genome. *S. scombrus* transcriptome, and the *S. japonicus De Novo* transcriptome. DN = *De Novo*, RG = Reference Genome, RT= Reference Transcriptome. To quantify transcripts, the salmon estimation method was used for the De Novo transcriptome and reference transcriptome. HiSat2 was used to map individual reads from each sample back to the reference genome.

|  | Raw Read Totals |  |  | Adapter Removal 1 |  |  | Adapter Removal 2 |  |  | QC Phred 30 |  |  | PolyA Tail Removal |  |  | Normilzation @ 30 |  |  |
| --- | --- | --- | --- | --- | --- | --- | --- | --- | --- | --- | --- | --- | --- | --- | --- | --- | --- | --- |
| Samples | Duplicate | Unique | Total | Duplicate | Unique | Total | Duplicate | Unique | Total | Duplicate | Unique | Total | Duplicate | Unique | Total | Duplicate | Unique | Total |
| MBA-SF-03_R1 | 8360164 | 38404707 | 46764871 | 7152686 | 34573750 | 41726436 | 7156543 | 34564307 | 41720850 | 10231787 | 29123982 | 39355769 | 10227822 | 29056712 | 39284534 | 6267066 | 7125268 | 13392334 |
| MBA-SF-03_R2 | 12982405 | 33782466 | 46764871 | 11750629 | 29975807 | 41726436 | 11755599 | 29965251 | 41720850 | 18334391 | 21021378 | 39355769 | 18325016 | 20959518 | 39284534 | 9617986 | 3774348 | 13392334 |
| MBA-SF-10_R1 | 6346944 | 43452896 | 49799840 | 5529860 | 26810923 | 32340783 | 5530842 | 26808515 | 32339357 | 7276707 | 22050125 | 29326832 | 7270962 | 21995021 | 29265983 | 5136459 | 6910384 | 12046843 |
| MBA-SF-10_R2 | 13356199 | 36443641 | 49799840 | 9817625 | 22523158 | 32340783 | 9819991 | 22519366 | 32339357 | 14894292 | 14432540 | 29326832 | 14884748 | 14381235 | 29265983 | 9052037 | 2994806 | 12046843 |
| MBA-SF-11_R1 | 7447013 | 27278799 | 34725812 | 8414269 | 31628023 | 40042292 | 8417197 | 31623787 | 40040984 | 10749140 | 27201207 | 37950347 | 10739227 | 27183031 | 37922258 | 6594493 | 5098173 | 11692666 |
| MBA-SF-11_R2 | 10707052 | 24018760 | 34725812 | 12015609 | 28026683 | 40042292 | 12017956 | 28023028 | 40040984 | 17347826 | 20602521 | 37950347 | 17338913 | 20583345 | 37922258 | 8949422 | 2743244 | 11692666 |
| MBA-SF-07_R1 | 6865542 | 30905326 | 37770868 | 7905694 | 33963600 | 41869294 | 7909000 | 33958788 | 41867788 | 10088353 | 29087070 | 39175423 | 10076734 | 29072832 | 39149566 | 6087065 | 4657799 | 10744864 |
| MBA-SF-07_R2 | 11419437 | 26351431 | 37770868 | 11767026 | 30102268 | 41869294 | 11767372 | 30100416 | 41867788 | 16420889 | 22754534 | 39175423 | 16416474 | 22733092 | 39149566 | 8263252 | 2481612 | 10744864 |
| MBA-SF-08_R1 | 10576915 | 38304010 | 48880925 | 5498657 | 41156009 | 46654666 | 5500736 | 41151844 | 46652580 | 8476065 | 34246713 | 42722778 | 8473448 | 34217083 | 42690531 | 4833483 | 8191452 | 13024935 |
| MBA-SF-08_R2 | 14404102 | 34476823 | 48880925 | 12434178 | 34220488 | 46654666 | 12426799 | 34225781 | 46652580 | 19785708 | 22937070 | 42722778 | 19780323 | 22910208 | 42690531 | 9310953 | 3713982 | 13024935 |
| MBA-SF-09_R1 | 9138629 | 37527577 | 46666206 | 6358501 | 24314827 | 30673328 | 6360078 | 24311954 | 30672032 | 8233105 | 20315485 | 28548590 | 8229024 | 20303561 | 28532585 | 4977404 | 4334833 | 9312237 |
| MBA-SF-09_R2 | 13085018 | 33581188 | 46666206 | 9618602 | 21054726 | 30673328 | 9618939 | 21053093 | 30672032 | 13986466 | 14562124 | 28548590 | 13983258 | 14549327 | 28532585 | 7082138 | 2230099 | 9312237 |
| MBA-WT-01_R1 | 6022403 | 36453616 | 42476019 | 4591287 | 28642545 | 33233832 | 4591413 | 28638842 | 33230255 | 7117676 | 24460458 | 31578134 | 7112342 | 24410151 | 31522493 | 4653011 | 7683023 | 12336034 |
| MBA-WT-01_R2 | 9708594 | 32767425 | 42476019 | 7903018 | 25330814 | 33233832 | 7903625 | 25326630 | 33230255 | 13692544 | 17885590 | 31578134 | 13687695 | 17834798 | 31522493 | 8143099 | 4192935 | 12336034 |
| MBA-WT-02_R1 | 8181458 | 20563794 | 28745252 | 5874877 | 14881066 | 20755943 | 5870704 | 14882987 | 20753691 | 7036712 | 12078843 | 19115555 | 7033544 | 12010327 | 19043871 | 5407129 | 4688477 | 10095606 |
| MBA-WT-02_R2 | 10935112 | 17810140 | 28745252 | 8305010 | 12450933 | 20755943 | 8305967 | 12447724 | 20753691 | 11286403 | 7829152 | 19115555 | 11282621 | 7761250 | 19043871 | 7872489 | 2223117 | 10095606 |
| MBA-WT-04_R1 | 9957661 | 31271583 | 41229244 | 7398159 | 23296559 | 30694718 | 7393715 | 23296755 | 30690470 | 9461042 | 19680097 | 29141139 | 9456992 | 19645312 | 29102304 | 6460112 | 5419375 | 11879487 |
| MBA-WT-04_R2 | 12924480 | 28304764 | 41229244 | 10039946 | 20654772 | 30694718 | 10035645 | 20654825 | 30690470 | 14316404 | 14824735 | 29141139 | 14314464 | 14787840 | 29102304 | 8818760 | 3060727 | 11879487 |

|  |  | DESeq2 | TransDecoder | KEGG |
| --- | --- | --- | --- | --- |
| 1 | Asymt_Wild | 2,155 | 806 | 567 |
| 2 | Sympt_Wild | 4,183 | 793 | 526 |
| 3 | Asympt_Sympt | 85 | 35 | 22 |

